# DNA barcoding revealed a high percentage of mislabeling in commercial fish products: the first empirical survey in South Texas

**DOI:** 10.1101/2023.08.29.555441

**Authors:** Rita B. Tommasi, Sanjana M. Lamia, Dysocheata Van, Isaiah Estrada, Zoen L. Kuchler, Dean Ramsey, Jyoti Tamang, Gen Kaneko, Hashimul Ehsan

## Abstract

Seafood mislabeling is a widespread problem that have produced a growing distrust of seafood industry. In this study, we examined the prevalence of mislabeling in fish samples from regional markets in the greater Houston area and close-by coastal communities. A total of 63 fish fillet samples were purchased, labeled, and stored at -20°C in individual packages until DNA extraction. DNA fragments of roughly 700 base pairs encoding cytochrome oxidase I (COI) were PCR-amplified from each DNA sample. With 99.6–100% nucleotide identity in the GenBank and BOLD databases, all samples were correctly identified at the species level. The scientific names identified by DNA barcoding were matched with legally acceptable market names using the US Food and Drug Administration (FDA) Seafood List. Out of the 63 samples examined, 13 samples (20.6%) were mislabeled. Moreover, 24 samples (38.1%) did not use the acceptable market names, indicating that the FDA policy is poorly implemented in seafood industry. The first DNA barcoding survey this area warranted the need of continuous monitoring and the dissemination of the regulation combined with taxonomic knowledge.

## Introduction

Fish mislabeling is a typical form of economic deception for the consumer. Intentional mislabeling is outright fraudulent, in which cheap fish is sold with a name of expensive fish such as halibut, swordfish, and snapper (Rasmussen & Morrissey, 2009). The confusion between the common name, acceptable market name, and scientific name is another source of fish mislabeling. For example, “flounder” includes 49 and 68 different species in FishBase and the US Food and Drug Administration (FDA) Seafood List, respectively (Food and Drug Administration, 2018; Froese & Pauly, 2023). To avoid misleading consumers, the European Union requires declaration of the scientific name for unprocessed products to ensure robust seafood labeling (Paolacci et al., 2021). However, the US remains lenient, originating about 40% of the world’s mislabeled seafood (Kroetz et al., 2020). The regular assessment of fish product authenticity is of a particular importance to ensure the quality and food safety in the US fish market.

DNA barcoding has been a useful tool in identifying species. Mitochondrial DNA (mtDNA) is a major target in DNA barcoding because: (1) it generally lacks large noncoding regions; (2) it is abundant and relatively easy to analyze; (3) it does not undergo genetic rearrangements such as recombination; (4) sequence ambiguities resulting from heterozygous genotypes can be avoided, and (5) regions with low intraspecies and high interspecies variation are already identified. Although several mtDNA regions such as 16S rRNA (Itoi et al., 2005; Kochzius et al., 2008) and cytochrome b (Ha et al., 2018; Itoi et al., 2020; Pepe et al., 2005) have been used for DNA barcoding, the 5’ region of cytochrome c oxidase subunit I (COI) has become a dominant region for identifying fish mislabeling (Di Pinto et al., 2013; Panprommin et al., 2020).

With a coastal extension of 450 miles, Texas is home to a large portion of fish commerce. Near-shore wetlands provide efficient natural water filters through plants and soils, serving as important nurseries for fish, crab, and shellfish in The Gulf of Mexico. A total economic value of commercial fisheries landed in Texas exceeded $168 million in 2022 (NOAA, 2022), and correct labeling is an important obligation to retain ecological and financial responsibility of the state. However, we did not find any DNA barcoding studies on seafood mislabeling specifically targeting local markets or restaurants in Texas — only two previous studies analyzed fish samples from this region to our knowledge. A previous DNA barcoding study included a sushi restaurant in Austin along with those in New York and San Francisco as cities where consumers likely demand high-quality food (Khaksar et al., 2015). Forty-three fish samples from Austin and Houston were analyzed in 2012 as a part of a nation-wide fish mislabeling survey (Warner et al., 2019). While the public inspection should be in effect, considering the scale and period of these previous studies, a new DNA barcoding survey, even preliminary, will bring new insights into how fish products are handled in this large economic hub of fish commerce.

In the present study, we tested fish samples from local markets in the greater Houston area and nearby coastal cities. Several samples were mislabeled, but we also found a high number of unaccepted market names. Based on the results, possible future strategies to reduce fish mislabeling were discussed.

## Methods

### DNA barcoding

Fish fillet samples (n = 63) were purchased from local markets in South Texas coastal cities (Table 1, mainly from the Houston area). All samples were labeled and stored at -20°C in individual packages until DNA extraction. A 10 - 50 mg piece of fast muscle tissue was cut from each sample, being sure to avoid contamination. A part of the samples was then analyzed by a BioRad Fish Barcoding Kit (Bio-Rad Laboratories Inc., Hercules, CA) according to the manufacturer’s instructions. PCR condition was 2 min at 94°C, followed by 35 cycles of 30 sec at 94°C, 2 min at 55°C, 1 min at 72°C. The final extension step was 10 min at 72°C. The PCR products were purified by a NucleoSpin Gel and PCR Clean-up kit (Macherey-Nagel., Düren, Germany). COI gene fragments were amplified from some samples according to the USDA recommended protocol (https://www.fsis.usda.gov/sites/default/files/media_file/2020-11/CLG-FPCR.pdf) using two primers: FishCO1LBC_m13F (5’-CACGACGTTGTAAAACGACTCAACYAATCAYAAAGATATYGGCAC-3’) and FishCO1HBC_m13R (5’-GATAACAATTTCACACAGGACTTCYGGGTGRCCRAARAATCA-3’).

**Table 1.** Fish samples analyzed in the present study.

| Table 1. Fish samples analyzed in the present study |  |  |  |  |  |  |
| --- | --- | --- | --- | --- | --- | --- |
| Label | Possible species | BOLD/BLAST top hit | BOLD/BLAST identity (%) | Mislabeled | Accession No. | Length |
| Pollock | See Table S1 | *Gadus chalcogrammus* | 99.83 | No | MH119970 | 609 |
| Pacific cod* | *Gadus macrocephalus* | *G. chalcogrammus* | 100 | **Yes** | MK283727 | 508 |
| Pacific cod* | *Gadus macrocephalus* | *G. macrocephalus* | 100 | No | MK283728 | 585 |
| Pacific cod* | *Gadus macrocephalus* | *G. macrocephalus* | 100 | No | MK283729 | 587 |
| Pacific cod* | *Gadus macrocephalus* | *G. macrocephalus* | 100 | No | MK283730 | 608 |
| Pacific cod* | *Gadus macrocephalus* | *G. macrocephalus* | 100 | No | MK283731 | 585 |
| Alaskan cod* | Considered as Alaska cod *Gadus macrocephalus* | *G. macrocephalus* | 100 | No | MK283715 | 593 |
| Alaskan cod* | Considered as Alaska cod *Gadus macrocephalus* | *G. macrocephalus* | 100 | No | MK283718 | 627 |
| Atlantic cod* | *Gadus morhua* | *G. morhua* | 99.83 | No | OR461757 | 581 |
| Pacific whiting | *Merluccius productus* | *M. productus* | 100 | No | MK283720 | 627 |
| Pacific whiting | *Merluccius productus* | *M. productus/angustimanus* | 100 | No | MK283732 | 617 |
| Pacific whiting | *Merluccius productus* | *M. productus/angustimanus* | 100 | No | MK283733 | 596 |
| Pacific whiting | *Merluccius productus* | *M. productus/angustimanus* | 100 | No | MK283734 | 610 |
| Pacific whiting | *Merluccius productus* | *M. productus/angustimanus* | 100 | No | MK283735 | 610 |
| Pacific whiting | *Merluccius productus* | *M. productus/angustimanus* | 100 | No | MK283736 | 597 |
| Atlantic croaker* | *Micropogonias undulatus* | *M. undulatus* | 100 | No | KX163997 | 670 |
| Yellow croaker* | *Larimichthys polyactis* | *Barbonymus altus* | 100 | **Yes** | MH119965 | 651 |
| Flounder | See Table S1 | *Lepidopsetta polyxystra* | 100 | No | MK283711 | 589 |
| Flounder | See Table S1 | *Lepidopsetta polyxystra* | 100 | No | MK283716 | 618 |
| Flounder | See Table S1 | *Lepidopsetta polyxystra* | 100 | No | MK283717 | 610 |
| Flounder | See Table S1 | *Limanda aspera* | 100 | No | MK283721 | 587 |
| Flounder | See Table S1 | *Limanda aspera* | 100 | No | MK283722 | 598 |
| Flounder | See Table S1 | *Limanda aspera* | 100 | No | MK283723 | 610 |
| Flounder | See Table S1 | *Limanda aspera* | 100 | No | MK283724 | 588 |
| Flounder | See Table S1 | *Limanda aspera* | 100 | No | MK283725 | 632 |
| Flounder | See Table S1 | *Limanda aspera* | 100 | No | MK283726 | 593 |
| Summer flounder* | *Paralichthys dentatus* | *P. dentatus* | 100 | No | KX164002 | 629 |
| Atlantic halibut* | *Hippoglossus hippoglossus* | *H. stenolepis* | 100 | **Yes** | KX164003 | 653 |
| Halibut | *Hippoglossus hippoglossus* or *H. stenolepis* | *H. stenolepis* | 100 | No | MK283719 | 649 |
| Monkfish | See Table S1 | *L. americanus* | 100 | No | MH760553 | 563 |
| Monkfish | See Table S1 | *L. americanus* | 100 | No | MH760554 | 579 |
| Monkfish | See Table S1 | *L. americanus* | 100 | No | MH760555 | 589 |
| Monkfish | See Table S1 | *L. americanus* | 100 | No | MH760556 | 496 |
| Monkfish | See Table S1 | *L. americanus* | 100 | No | MH760557 | 432 |
| Yellowfin tuna* | *Thunnus albacares* | *Xiphias gladius* | 100 | **Yes** | MH119957 | 597 |
| Yellowfin tuna* | *Thunnus albacares* | *T. albacares* | 100 | No | MH119959 | 334 |
| Amberjack | See Table S1 | *Seriola dumerili* | 100 | No | MH119972 | 669 |
| Blue runner | *Caranx crysos* | *C. crysos* | 100 | No | MH119976 | 633 |
| Mackerel | See Table S1 | *Atule mate* | 99.62 | **Yes** | MH119973 | 588 |
| King mackerel | *Scomberomorus cavalla* | *S. cavalla* | 100 | No | MH119964 | 663 |
| Norway mackerel* | Considered as mackerel; see Table S1 | *Scomber scombrus* | 100 | No | MH119971 | 669 |
| Atlantic pomfret* | *Brama brama* | *Peprilus medius* | 100 | **Yes** | MH119960 | 669 |
| Big grouper* | Considered as giant grouper *Epinephelus fucoglottus* x *E. lanceolatus* | *Hyporthodus niphobles* | 100 | **Yes** | MH119963 | 462 |
| Black drum* | *Pogonias cromis* | *Pogonias cromis* | 100 | No | KX164000 | 605 |
| Seabass | See Table S1 | *Dissostichus eleginoides* | 100 | **Yes** | MH119956 | 480 |
| Snapper | See Table S1 | *Pristipomoides multidens* | 100 | No | MH119958 | 437 |
| Round scad* | *Decapterus punctatus* | *D. macarellus* | 99.85 | **Yes** | MH119969 | 675 |
| Round scad* | *Decapterus punctatus* | *D. macarellus* | 100 | **Yes** | MH119978 | 663 |
| Starry butterflyfish* | Considered as butterflyfish; see Table S1 | *Peprilus medius* | 100 | **Yes** | MH119975 | 666 |
| Shark | See Table S1 | *Carcharhinus brevipinna* | 100 | No | MH119961 | 675 |
| Striped Pangasius (Swai) | *Pangasianodon hypophthalmus* | *Pristipomoides multidens* | 100 | **Yes** | MK283712 | 640 |
| Swai | *Pangasinodon hypothalamus* | *P. hypophthalmus* | 100 | No | MH119968 | 222 |
| Farmed Swai | *Pangasinodon hypothalamus* | *P. hypophthalmus* | 100 | No | MH119967 | 639 |
| Natural Tilapia* | Considered as tilapia; see Table S1 | *Oreochromis niloticus* | 100 | No | MK283714 | 560 |
| Catfish | See Table S1 | *Ictalurus punctatus/dugessii* | 100 | No | MK283713 | 624 |
| Catla | *Gibelion catla (Labeo catla)* | *G. catla* | 100 | No | KX163998 | 629 |
| Grass carp | *Ctenopharyngodon idella* | *C. idella* | 100 | No | MH119977 | 615 |
| Rohu | See Table S1 | *Labeo rohita* | 100 | No | KX164001 | 669 |
| White perch | *Morone americana* | *M. americana* | 100 | No | MH119962 | 675 |
| White perch | *Morone americana* | *M. americana* | 100 | No | MH119974 | 606 |
| Ganges river sprat* | *Corica soborna* | *C. soborna* | 100 | No | KX164004 | 647 |
| Gangetic ailia | *Ailia coila* | *A. coila* | 100 | No | KX163999 | 658 |
| River barb* | Considered as barb; see Table S1 | *Micropogonias undulatus* | 100 | **Yes** | MH119966 | 660 |

PCR was conducted at 95°C for 3 min, followed by 34 cycles of 95°C for 30 s, 55°C for 30 s, and 72°C for 1 min. The final extension step was at 72°C for 5 min. DNA sequences were determined by Eurofins Genomics (Louisville, KY). The obtained sequences were analyzed by the Barcode of Life Data (BOLD) system (http://www.boldsystems.org/) to determine species. The BLASTN program at the NCBI BLAST (https://blast.ncbi.nlm.nih.gov/Blast.cgi) was used with default parameters when the determined COI sequence was shorter than 500 bp. US Food and Drug Administration Seafood List (Food and Drug Administration, 2018) and FishBase (Froese & Pauly, 2023) were used to retrieve a list of possible species from labels (Supplementary Table S1).

## Results and Discussion

A DNA barcoding survey on fish mislabeling can choose one of the two strategies: testing a wide variety of species (Lakra et al., 2016; Pardo et al., 2018; Smith et al., 2008) or focusing on a specific group of fish (Bektas et al., 2019; Di Pinto et al., 2013; Lowenstein et al., 2009). We employed the former strategy since the present study was the first DNA barcoding survey specific to this geographical area to our knowledge. However, we included relatively higher numbers of expensive, white-fleshed species such as cod, flounder, and monkfish, which tend to be mislabeled (Hu et al., 2018).

The COI gene sequences solely containing the coding region were successfully determined for the 63 samples using the conventional PCR sequencing approach (Table 1). The average length of COI region determined was 595.5 bp. All samples were identified at the species level with 99.62-100% nucleotide identity. Out of the 63 samples, 13 samples (20.6%) were determined to be mislabeled, which was generally comparable to the lower end of mislabeling frequencies reported in US and Mexico: 15-17% for grocery in a nation-wide survey (Warner et al., 2019), 16.3% in restaurants in Austin, New York, and San Francisco (Khaksar et al., 2015). Higher mislabeling rates have also been reported such as 30.8% in markets and restaurants in Mazatlan, Mexico City, and Cancun (Munguia-Vega et al., 2022) and 47% in sushi restaurants in Los Angeles (Willette et al., 2017). Our previous survey on the genetically modified (GM) plants have found no false labeling in organic products, which should not contain any GM plants (Tegeler et al., 2020). These results may indicate the overall honest attitude of food industries toward food labeling in this area. In parallel, the low mislabeling rate is attributed to the generous regulation in US labeling guideline (Food and Drug Administration, 2018). When FishBase was used to match the scientific name and common name, as commonly done in previous studies (Lamendin et al., 2015; Panprommin & Manosri, 2022), the mislabeling rate was 36.5% (23 out of 63, Supplementary Tables S2 and S3).

As shown in Table 1, 2 samples were mislabeled in the 17 white-fleshed marine fish (pollock, cod, Pacific whiting, and croaker). Pacific cod is a common name exclusively used for *Gadus macrocephalus* (FishBase, Supplementary Table S3), but this species was substituted with Alaska pollock *G. chalcogrammus*. These species share a similar morphology, and this substitution has also been reported in other studies (Feldmann et al., 2021; Helgoe et al., 2020). Moreover, many fish products did not have acceptable market names. For example, only “cod” and “Alaska cod” are acceptable market names of *G. macrocephalus* and *G. morhua* (FDA Seafood List, Supplementary Table S1), and thus “Pacific cod” and “Alaskan cod” samples have legal problems. Pacific whiting samples were identified as a correct species *Merluccius productus*, although the COI sequence also showed 100% nucleotide identity with Panama hake *M. angustimanus*. While this represents a limitation of DNA barcoding based on the COI gene, these two species are genetically close to each other. Both Atlantic croaker and yellow croaker must be labeled just as “croaker” (Supplementary Table S1). Yellow croaker *Larimichthys polyactis* was substituted with red tailed tinfoil *Barbonymus altus*, which belongs to the family Cyprinidae (freshwater fish such as carp and minnow). This substitution is likely an intentional fraudulent mislabeling.

We analyzed 12 flatfish samples including flounder and halibut. The flounder samples were identified as Northern rock sole *Lepidopsetta polyxystra* and yellowfin sole *Limanda aspera* (Table 1). These species are acceptable as “flounder” in the FDA Seafood list, but not in the FishBase. The difference accounted for the discrepancy in the mislabeling rates, 20.6% based on the FDA Seafood list and 36.5% based on FishBase. Summer flounder and Atlantic halibut were unacceptable market names.

Three species in the genus *Lophius* can be sold as monkfish in US (Supplementary Table S1). Our DNA barcoding showed that all monkfish samples were *L. americanus*, identified no mislabeling. The angel shark *Squatina squatina* used to be sold as monkfish, exposing the species to the risk of extinction (Porcher & Darvell, 2022), but this was not the case in the present study.

Other marine fish products showed relatively high frequencies of mislabeling and unaccepted market names (Table 1, from yellowfin tuna to shark). Among them, the substitution of yellowfin tuna *Thunnus albacares* with swordfish *Xiphias gladius* and that of seabass with Patagonian toothfish *Dissostichus eleginoides* could be intentional. The substituted species are relatively expensive and differ from the substituting species in morphology and distribution. Others may be attributed to the confusion. In this study, we analyzed a relatively high number of white-fleshed species that were commonly mislabeled (Hu et al., 2018). However, since the mislabeling frequency was higher than expected in other marine fish species, further studies could focus on these species with increased number of samples.

Two mislabeling cases were found in inexpensive freshwater fish (Table 1, swai to river barb). Swai *Pangasianodon hypophthalmus* was substituted with a marine fish, Goldbanded jobfish *Pristipomoides multidens*. This fish is also called Goldband snapper and is generally considered as a palatable fish species. River barb turned out to be Atlantic croaker *Micropogonias undulatus*, which is also a high-value species. While more samples are needed to reach a conclusion, these mislabeling cases may be attributed to confusion and/or poor traceability.

In summary, the first fish DNA barcoding study in the greater Houston area demonstrated that 20.6% of fish fillet samples were mislabeled, and 38.1% of the samples used market names. Several approaches that have been proposed in previous studies should facilitate the solution of this problem (Cawthorn et al., 2018; Hu et al., 2018). Namely, mandatory inclusion of scientific names is the first policy to be implemented, which would eliminate the ambiguity between the scientific name, common name, and acceptable market name. Inclusion of geographical origin and harvest methods also benefit consumer. Regular DNA barcoding survey well acquainted with local community will help consumers to make educated decisions as well as encourage the seafood industry to use more accurate labeling.

## Acknowledgements

The authors are grateful to the M.G. and Lillie A. Johnson Foundation, Victoria, Texas, for supporting students involved in this research.

